# Multi-channel high-density single-molecule localization

**DOI:** 10.64898/2026.07.24.740199

**Authors:** Hao Sha, Lucas-Raphael Müller, Nestor Miguel Castillo Duque de Estrada, Maxime Mathieu, Arthur Jaques, Zach Marin, Yongbing Zhang, Jakob H. Macke, Jonas Ries

## Abstract

Deep learning has enabled single-molecule localization microscopy (SMLM) at high emitter densities, but only for single channel systems. Here we present DECODE-Plex, a deep-learning-based framework to localize dense single molecules with overlapping point spread functions simultaneously in multiple channels. We showcase DECODE-Plex on experimental ultra-high density dual-color and 3D live-cell data. Packaged for ease of use, it will enable many groups to improve imaging speed and quality of multi-channel SMLM.

---

Single-molecule localization microscopy (SMLM) is a super-resolution method that can resolve cellular structures at nanometer resolution^1–3^. The basis of SMLM is to label structures of interest with switchable fluorophores and to acquire tens of thousands camera frames in which only a small fraction of the fluorophores is activated, to avoid overlapping emitters. The position of these sparse emitters is then localized using, e.g., maximum-likelihood fitting^4,5^. The need for sparsity and, consequently, many camera frames, leads to poor temporal resolution and low throughput and puts an upper bound on the labeling density. This major limitation of SMLM was reduced with deep learning algorithms like DeepSTORM^6–8^, DECODE^9^, its extensions^10–13^, and related approaches^14^, which allow for fitting fluorophores at high densities with overlapping point-spread functions (PSFs), outperforming even dedicated multi-emitter fitting algorithms^15^. However, none of these programs are compatible with multi-channel SMLM approaches. Simultaneous detection of fluorophores in multiple channels is the basis for several advanced SMLM techniques: Ratiometric (de-mixing) multicolor SMLM^16,17^ splits the emission of spectrally overlapping fluorophores with a dichroic mirror onto two cameras or regions on one camera and uses relative intensities of the single emitters to assign the color. Biplane^18^ or multi-plane^19^ detection enables 3D localization, even in combination with spectral de-mixing^20^. 4Pi SMLM^21,22^ uses interference recorded on 3 or 4 channels for greatly improved z localization, even in multicolor^23^.

Algorithms that fit multi-channel data individually per channel cannot extract the full information and hence do not reach the best possible localization precision, described by the Cramér–Rao bound (CRB)^5^ in the single-emitter regime. To overcome this limitation, we previously developed GlobLoc^5^, an approach that uses maximum likelihood estimation on all channels simultaneously to maximize precision. However, as a single-emitter fitter, it is not compatible with high emitter densities beyond the true single-molecule regime.

Here, we present DECODE-Plex, a new multi-channel, high-density fitting algorithm based on deep learning. It builds on the architecture of DECODE^9^ and introduces a new representation for input and output that allows it to combine information across multiple input channels. It performs simultaneous detection and localization and consistently achieves better detection performance and localization precision compared to GlobLoc for biplane and multicolor SMLM.

For each frame (**Supplementary Figure 1**), the DECODE-Plex network outputs a detection probability, positions in x/y/z using offset vectors with respect to the pixel center, background estimates, and a photon count prediction per channel for multicolor data or a total photon count for biplane data (**Figure 1A**). An uncertainty estimate accompanies each output predictor. For multicolor SMLM, we use the predicted photon counts in both channels together with their uncertainties and the expected intensity ratio between the channels for a probabilistic assignment of the emitters’ colors (**Methods**). Like DECODE, DECODE-Plex uses temporal context, i.e., it considers that the same fluorophore can be visible in several adjacent camera frames.

**Fig. 1:**
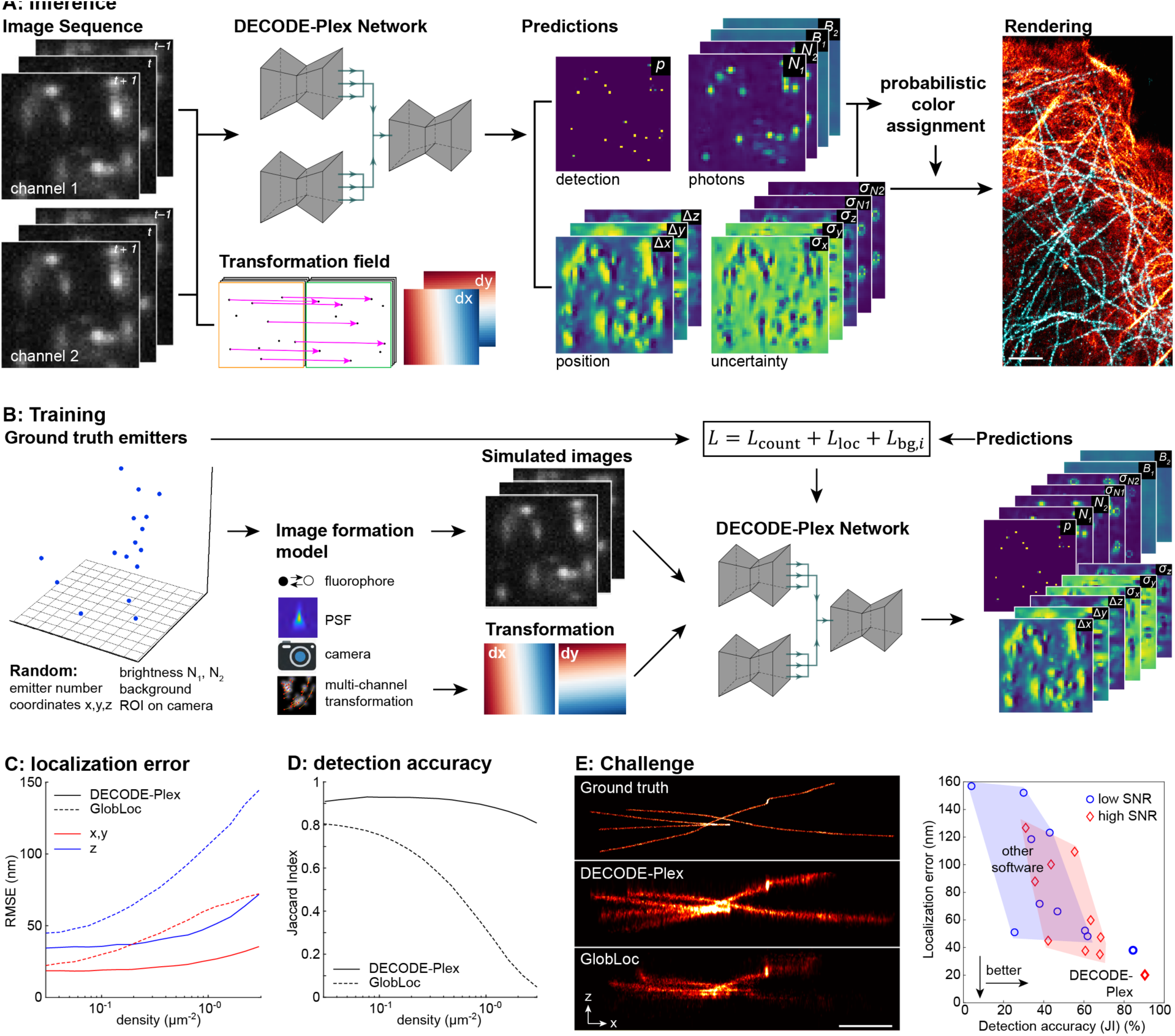
DECODE-Plex for high-density single-molecule localization, architecture and performance on simulated data. A: DECODE-Plex architecture. The neural network uses information from multiple channels simultaneously and is informed about their spatial relationships via the transformation field that denotes the pixel-wise offsets between the channels. The network maps its predictions onto frames representing a probability for detection, the photon count in each channel, positions, uncertainties related to these predictions, and a background. For biplane SMLM, the photon counts are linked. For multicolor SMLM, the photon counts in each channel are predicted and color assignment is performed by likelihood estimation. **B: Training.** DECODE-Plex is trained with simulated data. Emitters are sampled randomly over the field of view. In addition, a position on the camera and a corresponding transformation is sampled to cover all possible positions at inference time. **C: Localization error** in the lateral and axial direction in dependence on the emitter density. **D: Detection accuracy** (Jaccard index). See **Supplementary Table 1** for training parameters. **E: SMLM Challenge**, side view reconstruction of the training data set MT0.N2.HD-BP (see **Supplementary Figure 8**, scale bar 1 µm) and performance of DECODE-Plex in comparison to other software (test dataset MT4.N2.HD, evaluation by the challenge organizers).

In general, the channels might not be perfectly aligned, but may exhibit shifts, rotations or different magnifications. Spatial transformations between the channels can be extracted from calibration measurements on beads, or directly from blinking fluorophores^24^. We provide this transformation per pixel as an input to the network during inference (**Figure 1A**). This allows us to re-use a trained model even if we change the region of interest (ROI) on the camera or if the transformation changes.

We train the DECODE-Plex network with entirely simulated data (**Figure 1B**). To avoid structural bias, we simulate emitters at random 3D positions and calculate realistic simulated camera frames in all channels using a precise, experimentally derived PSF model^24^, a fluorophore blinking model, experimental distributions of photon counts and background values, a camera noise model, and random spatial offsets sampled from the transformation. During training, the negative log-likelihood loss, which describes how well the predictions explain the ground-truth data, is minimized (**Methods**).

The accuracy of the PSF model and transformation are of utmost importance, as a model mismatch can lead to a bias in the position output towards the pixel which appears as a pixelation artifact (**Supplementary Figure 2**). To obtain accurate PSF models, we use either the GlobLoc PSF calibration routine^5^ or uiPSF^24^, a framework that can learn multi-channel PSFs including transformations from experimental bead stacks or directly from experimental SMLM data. If needed, residual pixelation artifacts can be corrected by a debiasing procedure (**Supplementary Figure 3**).

We first evaluated the performance of DECODE-Plex on simulated data using metrics established by the SMLM Challenge^15^, which is a benchmarking resource to aid users in the choice of localization algorithms. We extended the challenge’s metrics for use with multicolor data (see **Methods**) and report the color assignment accuracy and rejection rate, i.e., the fraction of correctly assigned colors and the fraction of rejected color assignments.

On simulated, single-frame, low-density data, DECODE-Plex performance approaches the Cramér–Rao bound (**Supplementary Figure 4**), i.e., the best theoretically possible precision, and thus performs equally as well as GlobLoc^5^. This demonstrates that the DECODE-Plex network can efficiently combine information across channels. For higher densities, DECODE-Plex outperforms GlobLoc for all simulated multicolor and biplane data in all metrics (**Figure 1C**, **Supplementary Figure 5**). In the SMLM Challenge^15^ on the dual-channel biplane test data sets, DECODE-Plex outperformed all other algorithms by a large margin (**Figure 1E**).

Next, we evaluated DECODE-Plex on experimental data. We imaged microtubules together with actin (**Figure 2A**) and the endoplasmic reticulum (**Figure 2B**) at localization densities about one order of magnitude higher than what we use for standard, single-emitter density SMLM. We used a 3D, astigmatic, dual-channel setup^25^ and labeled each pair of proteins with two far-red dyes with highly overlapping emission spectra. For these experimental high-density datasets, we consistently find better results with DECODE-Plex than with GlobLoc. In particular, regions with high local density were better resolved with DECODE-Plex, especially in the axial direction. In addition, DECODE-Plex retains more localizations because GlobLoc filters out localizations with overlapping PSFs based on the log-likelihood of the fit.

**Fig. 2:**
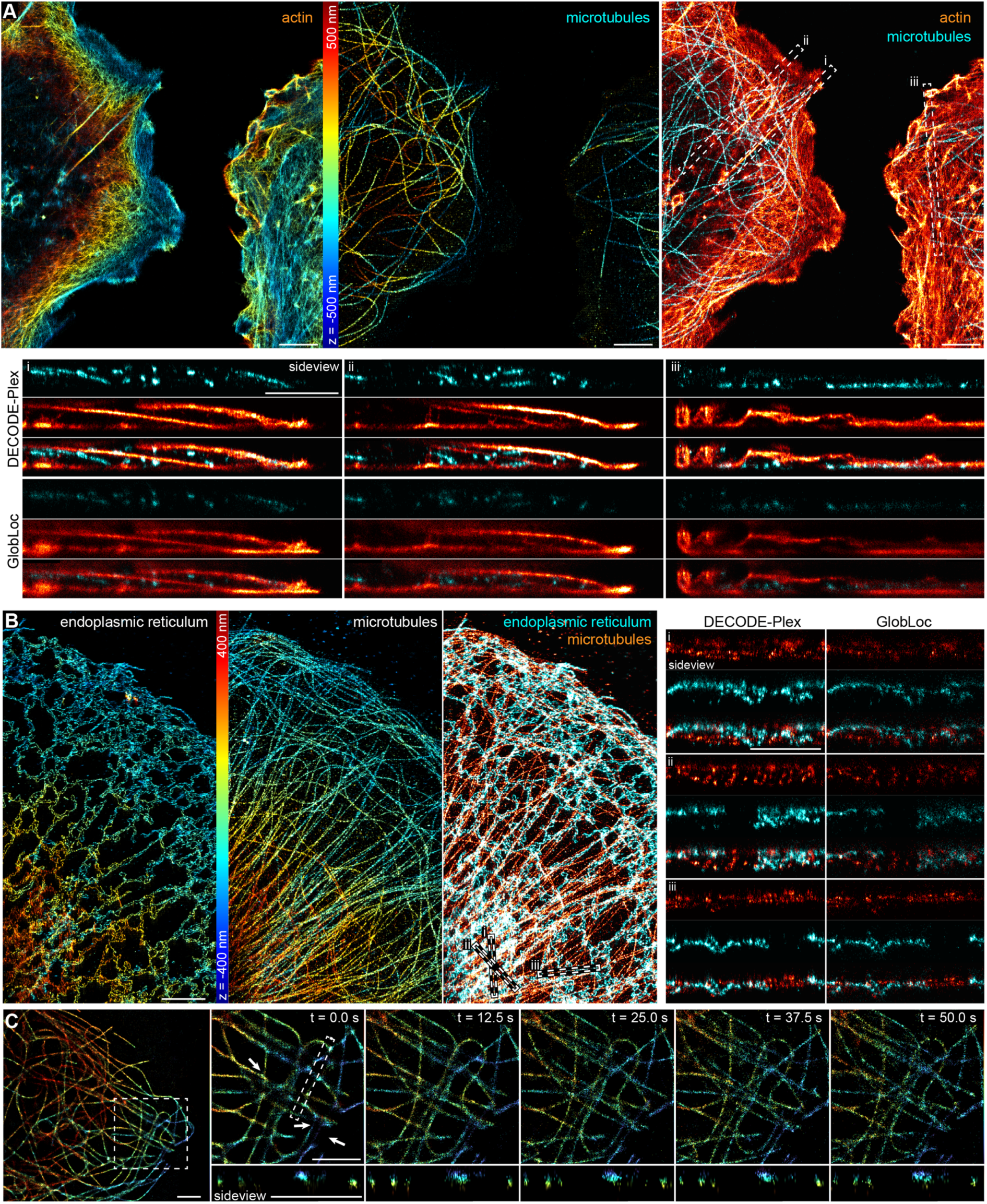
Performance on experimental high-density data. **A: Dual-color SMLM on actin** (AF647, phalloidin+) **and microtubules** (α- and β-tubulin, CF660C), comparing DECODE-Plex and GlobLoc. Side view reconstructions as indicated in the overlay image. **B: endoplasmic reticulum (**Nogo-B, AF647**) and microtubules (**α- and β-tubulin, CF660C**). C: Live-cell 3D SMLM on microtubules** using biplane imaging and the self-blinking fluorophore HMSiR-tubulin^27^. Scale bars 2 µm.

Compared to measurements at single-emitter densities, DECODE-Plex reduces the imaging time by around one order of magnitude. These high imaging speeds are critical for live-cell measurements, where the time resolution needs to match the biological process under investigation. We demonstrated the performance of DECODE-Plex by imaging microtubules in living cells in 3D with the biplane approach. Indeed, we could observe rearrangements of the microtubule cytoskeleton on the nanometer scale in 12.5 s intervals. The higher localization density and faster imaging speeds of DECODE-Plex are achieved without increasing the excitation laser power and light dose. This is important, as even in our measurements, phototoxicity was the limiting factor, perturbing the microtubule organization after approximately 2 minutes of imaging.

The DECODE-Plex network needs to be retrained whenever the PSF changes. Using a warm start based on a previous training, this takes approximately 30 minutes (**Supplementary Figure 6**). A new training takes approximately 4 hours on an RTX4090 workstation (**Supplementary Figure 7**). Inference is usually faster than imaging (>60 frames per second), allowing for real-time analysis, and in contrast to GlobLoc it is independent of the number of fluorophores.

The innovations of DECODE-Plex that enable multi-channel inference can directly extend approaches that are based on DECODE^10–13,26^ to multiple channels and will likely be applicable to other neural network single-molecule localization approaches^6–8,14^. Accompanied by detailed documentation and tutorials, DECODE-Plex will enable a large and growing community to perform multi-channel, high-density SMLM, greatly increasing throughput and imaging speed, and opening new sample regimes.

## Methods

### DECODE-Plex network architecture

Our architecture builds upon a combination of modified 2D U-Net^28^ models as the neural network’s backbone (**Supplementary Figure 1**). In contrast to the original implementation, we use an exponential linear unit (ELU) activation function^29^ to improve training stability.

DECODE-Plex accepts multi-channel images of blinking fluorophores in a sliding temporal window as input (e.g. for a two-channel imaging setup and a three-frame temporal window, each network would process three consecutive frames at a time). Each channel is processed in its own U-Net (first feature extraction modules, **Supplementary Figure 1**), and the outputs of these single-channel U-Nets are fed to a final U-Net (feature extraction module + output module, **Supplementary Figure 1**) that combines this information with prior knowledge of the transformation between the channels. The spatial transformation between channels is an additional input that is made available to all U-Nets, and it is a passed as a collection of offset maps between the two channels (for a two-channel setup, these are two maps with the difference in position between channel 1 and channel 0 in the x-direction and y-direction, respectively).

Our network aims to predict (1) a single, combined probability map *p_k_* of having found an emitter (following a Bernoulli distribution) in near proximity to pixel *k* in all channels, (2) the parameter vector ***μ****_k_* = (*x_k_*, *y_k_*, *z_k_*, *N_1,k_*, *N_2,k_*, …) containing the three-dimensional displacements with respect to the center of each pixel and the photon counts *N_c,k_* for each channel *c*, the respective uncertainties **Σ***_k_* = (*σ_x,k_*, *σ_y,k_*, *σ_z,k_*, *σ_N_*_1,*k*_, *σ_N_*_2,*k*_, …), and a background prediction *b_c,k_* for each channel. The predicted positions are always reported with respect to one reference coordinate system, which we choose as, without loss of generality, that of the first channel.

### Loss function for multi-channel detection, localization, and uncertainty estimates

We built upon DECODE’s^9^ loss function as a combination of count loss *L_count_*, localization loss *L_loc_*and background loss *L_bg_*:

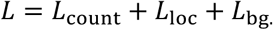

The count loss constrains the predicted number of emitters in each frame based on the network-predicted detection probability *p_k_* at pixel *k*:

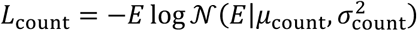

where *E* is the number of simulated emitters in the frame and N(.) is a normal distribution, 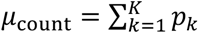, and 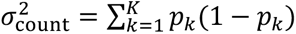, and *K* is the total number of pixels. As the network outputs a single, combined probability map for all channels, this loss is unchanged from the single channel DECODE approach.

The background loss penalizes deviations between the predicted and simulated background images. For multi-channel output, the photon count in the localization and background loss is extended to multiple channels. In the biplane mode, we link the photon count (*N_1,e_* = *r N_e_*, *N_2,e_* = (1 − *r*) *N_e_*), where the ratio *r* = *N_1,e_*/(*N_1,e_* + *N_2,e_*) is fixed to a value determined by the dual-channel PSF calibration. We still use a multi-channel background prediction and, therefore, multi-channel background loss:

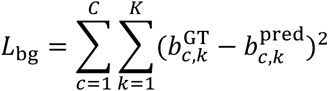

where *C* is the number of channels, 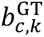 is the true background at pixel *k* in channel *c* and 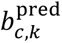 is the network-predicted value.

The localization loss optimizes the predicted emitter parameters and their associated uncertainty estimates. For each simulated ground-truth emitter *e* with parameter vector 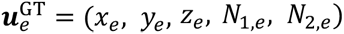 for multi-color data and 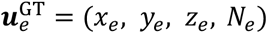 for biplane data, the network evaluates a mixture likelihood:

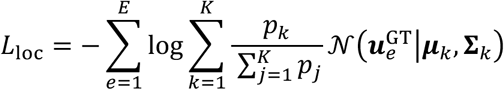

where ***μ****_k_* and **Σ***_k_* denote the corresponding predicted emitter parameters and uncertainty estimates, as described in the section **DECODE-Plex network architecture**. The ground truth parameter vector contains counts from all channels, thus adjusting the probability space over which the log likelihood is maximized. The rest of the loss function remains the same as in the single channel DECODE approach.

### Training details

Our model is trained purely on live simulated training data (see section **Training Data Simulation**). We perform training on 40 x 40 pixel-sized frames to give enough space for positional transformation between the channels. We sample the position on the camera to accommodate for the fact of varying positional transformations at different positions of the multi-camera setup (see section **Multi-channel transformation**). We use the AdamW optimizer^30^ with an initial learning rate of 1·10^−4^. The learning rate is reduced by a factor of 2 using the ReduceLROnPlateau scheduler whenever the validation loss plateaus^31,32^. We exclude dim emitters with less than 100 photons (summed over all channels) from the ground-truth emitter set used for the loss computation during training.

### Localization extraction and post-processing

#### Localization Extraction

Localization candidates are extracted from the frame representation by thresholding the probability output *p_k_*. Ideally, *p_k_* is sparse and well-separated into a bimodal distribution, and its sum corresponds to the true number of emitters. Small probability clusters can emerge under hard conditions instead of single sparse probability outputs. These local neighborhoods are integrated and form a cumulative probability threshold for localization extraction. The localization offsets and the uncertainty predictors are extracted at the thresholded pixels, and the localization position is computed as the pixel center positions plus the respective offsets.

#### De-Biasing

Under hard conditions, i.e., high density, low brightness or strongly out-of-focus fluorophores, the network tends to bias the localization estimates towards the center of the pixel (**Supplementary Figure 3**). The effect can be measured by computing the subpixel position histogram across the x and y dimensions. We allow for optional post-processing of these localizations to equalize the histogram. We observed the biasing to be correlated with the z position and, therefore, performed histogram equalization separately for different z ranges. Typically, z was divided into bins of 100 nm. Within each z bin, localizations were further grouped according to their predicted lateral localization uncertainty, and histogram equalization was performed independently for each uncertainty group.

#### Color Assignment

We perform the color assignment as a post-processing step. The emission wavelengths, dichroic beamsplitter characteristics and filters determine the (true) photon count ratio between the two channels. The expected ratio can be empirically measured or theoretically computed from the specification of the labels and optical components.

For each localized emitter, the predicted photon counts *N_c_* are normalized across all imaging channels to obtain the observed photon-ratio vector:

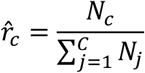

where *C* is the number of imaging channels. For each candidate color *c*, the expected photon-ratio vector is denoted by *r_c_*. Since DECODE-Plex outputs uncertainty measures for the photon counts *σ_Nc_*, we can compute the likelihood *L_c_* of the observed photon ratio vector under color *c* using a Gaussian model,

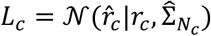

where *r_c_* is the expected photon-ratio vector for color *c*, N(⋅) denotes the Gaussian probability density function, the covariance matrix 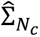 is constructed from the predicted photon count uncertainties normalized by the total photon count. The posterior probability of assigning an emitter to color *c* is then computed as:

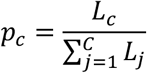

where *L_j_*is the likelihood evaluated for candidate color *j*.

### Training Data Simulation

We continuously simulate training data for DECODE-Plex on the fly and use each sample only once to avoid overfitting the U-Net backbones.

The performance of DECODE-Plex is tightly linked to the closeness of the simulation to the experimental data and the accuracy of the image formation model, which consists of several steps, as discussed in the following:

#### Point-Spread-Function

The PSF describes the image formed by the characteristics of the microscope and the object in the object plane. A single emitter at position ***r****_0_* = [*x_0_*, *y_0_*, *z_0_*] gives rise to an image *I*(*x*, *y*) = PSF(*x* − *x_0_*, *y* − *y_0_*, *z* − *z_0_*).

We use cubic spline interpolations as an accurate and flexible representation for arbitrary PSFs. We follow Li et al.^5^ and Babcock et al.^33^ and use our custom, optimized CUDA kernel, which allows us to simulate 100 000 frames in less than 0.5 s on an RTX 4090.

To obtain accurate image formation models for dual-channel simulation, we calibrated the optical system from experimentally acquired bead z-stacks using SMAP^5,34^ or uiPSF^24^. Notably, uiPSF also supports direct *in situ* PSF estimation from experimental single-molecule data, without requiring dedicated bead z-stack calibration. This calibration procedure provides channel-specific PSF models for each detection path, as well as the spatial transformation between channels required for consistent multi-channel simulations. This is necessary, as the choice of the center of the PSF is arbitrary for a spline-modelled PSF, thus PSF and transformation are not independent of one another.

The calibrated PSFs are subsequently represented by cubic spline interpolation and incorporated into the forward simulation of DECODE-Plex.

#### Camera

DECODE-Plex does not differentiate between a multi-camera setup and a multi-ROI, single-camera setup to model multiple channels. In all scenarios, we apply global, pixel-value preserving transformations to maximize the spatial overlap of all channels. This particularly includes de-mirroring and global shifts by multiples of pixels. The procedure ensures that the positional differences between the images of the same source emitter in the object plane are not too far apart for the final U-Net (see **Figure 1A, Supplementary Figure 1**) to combine all information^35^.

#### Shot noise and camera Noise

Following Sage et al.^15^, we model shot noise arising from the stochastic nature of photon detection by drawing a random Poisson variable for each pixel of the simulated images. In addition, for EMCCD cameras we model amplification noise from the electron multiplying gain by approximating it with a Gamma distribution, and we add readout noise approximated with a Gaussian distribution.

#### Multi-channel transformation

We assume the positional transformation in the multi-channel setting to follow an affine transformation *T*.

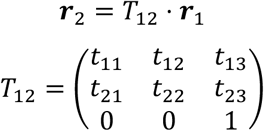

where ***r****_1_*= [*x_1_*, *y_1_*, 1] are the coordinates in the reference channel and ***r****_2_*= [*x_2_*, *y_2_*, 1] are the coordinates in a target channel.

*T_12_* is estimated jointly with the multi-channel PSF following the GlobLoc routine^5^ or with uiPSF^24^. Both the PSFs and the transformations can be extracted from a bead calibration or directly from the experimental data^24^.

At training time, we do not enforce any constraints on the relation between the photon counts in the respective channels to avoid biasing the photon predictions. Instead, we sample the photon count independently in multi-channel modes. The only exception is in the biplane mode, where only a single photon count is sampled and distributed across the channels according to the PSF’s normalization.

#### ROI Sampling

Since the frame size can vary between training and inference, we feed the transformation of the experimental data at inference time as an auxiliary frame for which we compute the displacement between the reference and the respective target channels in x and y for each pixel.

We sample the simulated camera position at training time to vary the ROI and displacement between the reference and respective channels (see section **Training details**).

#### Photophysics

We sample each fluorophore’s photon flux and initial appearance from a uniform distribution. With a uniform distribution, we argue that there is less risk of bias toward one of multiple channels in a multi-channel experiment which typically peak at different photon flux counts and have different lifetimes. In the case of a multi-channel experiment where the photon counts are unlinked, we sample the photon flux per channel independently. Furthermore, we draw the fluorophore ‘on time’ from an exponential distribution. The emitters are then distributed and discretized over the frames, which converts the photon flux by integration over time to a photon count. We do not model long-term re-appearance.

#### Estimating simulation parameters

Commonly, only the background and photon distribution will vary significantly between different SMLM experiments on a daily timescale. Covering the entire background regime is vital such that the model does not confuse background fluorescence with actual emitters. We found no penalty for training in a broad background regime (**Supplementary Figure 9**). We propose one of two options for regular usage of our algorithm: (A) Either train a broad distribution of photon and background and refine on data of particular interest or (B) perform a pre-fit with GlobLoc^5^ and use these parameters for training a specific DECODE-Plex model directly. With larger changes to the PSF (e.g., due to changes in the optical setup), new training is mandatory.

### Evaluation

We validated the performance of DECODE-Plex on simulated data using metrics established by the SMLM-Challenge^15^ and extended these metrics to multiple channels. We compared the performance to GlobLoc^5^, a state-of-the-art maximum likelihood (MLE) multi-channel fitter.

The Detection Accuracy describes how many true emitters are found and how many false localizations are reported. It is measured here as the Jaccard Index (JI), a relation between true positives (TP), false positives (FP), and false negatives (FN): JI = TP/(TP + FN + FP). True positives (TP) are determined by an assignment algorithm, a cost-minimizing assignment algorithm like the Hungarian algorithm^36^. Following the conventions established by the SMLM-Challenge, a predicted localization is considered a true positive if it can be matched to a ground-truth emitter within a lateral distance of 250 nm and an axial distance of 500 nm.

The localization precision (lateral, axial or volumetric) can be quantified for true-positive and matched positive pairs and is typically reported as the root mean squared error (RMSE):

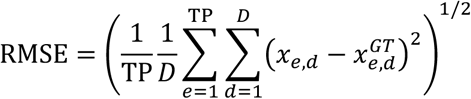

where *D* is the dimensionality, *x_d_* and 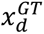 are the predicted and ground truth coordinates along their respective axes, respectively.

#### Multicolor Metrics

In addition to the metrics outlined above, the assignment of emitters to the correct color is vital for multicolor SMLM. Here, we report the Accuracy Acc, which describes the fraction of correct color assignments, and the Rejection rate Rej, which describes the fraction of unassigned emitters.

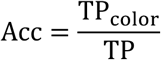

where TP*_color_*are the true positives with correct color assignment and TP are the true positives.

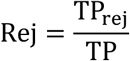

where TP*_rej_*are the true positives rejected due to fuzzy color assignment, and TP are the true positives.

#### Validation datasets

For multicolor data (**Supplementary Figure 5**), we used the following simulation parameters, comparable to our experiments: total photon flux of 5000 photons per emitter distributed across the two channels [ch0, ch1] according to channel-splitting ratios of [0.173, 0.827] and [0.019, 0.981], average on-times of [1.0, 3.0] frames and background count of [20, 135] photon in the two channels. We simulated a minimum of 10 000 emitters for each density step.

For biplane data (**Figure 1C**), we used: total photon flux of 10 000 photons per emitter equally distributed across the two channels, background of [50, 50] photons, and a lifetime average of 1.0 frame. We simulated a minimum of 10 000 emitters for each density step.

### Sample preparation and imaging

#### Seeding of U-2 OS cells on glass coverslips

Glass coverslips (Marienfeld, 117640) were treated overnight with a solution of 18.5% (v/v) hydrochloric acid in methanol with constant stirring, then washed thoroughly with deionized water and dried. Wild-type U-2 OS cells were seeded on the treated coverslips to reach between 30-50% confluence after 1-2 days of incubation at 37°C and 5% CO2 in DMEM 1x (Gibco, 11880036) supplemented with MEM NEAA 1x (Gibco, 12084947), GlutaMax 1x (Gibco, 35050038), 10% FBS (Gibco, 10270106) and ZellShield 1x (Minerva Biolabs, 13-0050).

#### Dual-color sample microtubules and actin

U-2OS cells were grown on glass coverslips, prepared as described before, and were then submitted to prefixation for 2 min using a solution of 0.3% (v/v) glutaraldehyde and 0.25% (v/v) Triton X-100 in Cytoskeleton Buffer (CB: 10 mM MES, pH 6.1, 150 mM NaCl, 5 mM EGTA, 5 mM glucose, 5 mM MgCl₂). Cells were then post-fixed for 10 min in 2% (v/v) glutaraldehyde in CB. Following fixation, samples were washed in PBS, permeabilized for 5 min with 0.2% (v/v) Triton X-100 in PBS, and blocked for 1 h with 2% (w/v) bovine serum albumin (BSA) in PBS. Cells were incubated overnight at 4°C with primary antibodies against α-tubulin [1:200] (Sigma, T6074) and β-tubulin [1:200] (Sigma, T5293) diluted in blocking solution. After three washes (5 min each) in PBS, samples were incubated for 2 h at room temperature with CF660C-conjugated anti-mouse IgG secondary antibody [1:500] (Biotium, 20815). Following three additional washes (5 min each), F-actin was labeled by incubation with Alexa Fluor 647 Plus phalloidin [1:3000] (Invitrogen, A22287) for 10 min. Coverslips were briefly rinsed in PBS and immediately mounted in blinking buffer (10 mM NaCl, 35 mM cysteamine, 10% (w/v) glucose, 500 µg/mL catalase, 40 µg/mL glucose oxidase in 50 mM Tris-HCl, pH 8.0) for dSTORM imaging.

#### Dual-color sample microtubules and endoplasmic reticulum

U-2 OS cells on glass coverslips were quickly rinsed with pre-warmed PEM buffer (80mM PIPES, 2mM MgCl2, 5mM EGTA, pH 6.8) and fixed with a solution of 4% paraformaldehyde, 0.1% glutaraldehyde, and 4% sucrose in PEM buffer for 10 min at 37°C. Then cells were rinsed in phosphate buffer (PB) (0.1 M sodium phosphate, pH 7.3) before quenched for 7 min at room temperature with a solution of 10mg/mL sodium borohydride in PB buffer. Next, cells were permeabilized and blocked for 1 hour with immunocytochemistry buffer (ICC buffer: 0.2% bovine serum albumin, 0.8% Triton X-100 in PB buffer), followed by 30 min incubation in a drop of Image-iT FX Signal Enhancer. Permeabilized cells were incubated overnight at 4°C with solution of antibodies against α-Tubulin [1:200] (Sigma, T6074), against β-Tubulin [1:200] (Sigma, T5293) and against Nogo-B [1:500] (R&D Systems, 18061422), all in ICC buffer, followed by three washes, 5 min each, in ICC buffer, and incubation with secondary antibodies CF660C-conjugated anti-mouse IgG [1:500] (Biotium, 20815) and AF647-conjugated anti-sheep IgG [1:500] (Thermo A21448), in ICC buffer, and finally washed with ICC buffer, three times, 5 min each, before mounting the sample for image acquisition in blinking buffer (10 mM NaCl, 35 mM cysteamine, 10% glucose, 500 g/L catalase from bovine liver, 40 g/L glucose oxidase, in 50 mM Tris/HCl pH 8.0).

#### Live-cell biplane sample

The living cells on coverslips were incubated for 1 hour in the same medium, with HMSiR-Tubulin^27^ added to a final concentration of 2 uM. Then, cells were briefly washed two times with fresh medium to reduce the background of free dye, and a new medium was finally added to the sample in the mounted chamber for imaging.

#### PSF calibration

TetraSpeck beads (Thermo Fisher Scientific, ref. T7279) were used to calibrate the PSF in both channels simulateneously. The glass coverslip was first activated by incubation with MgCl₂ for 3 min to promote adhesion of the beads to the glass surface. A bead suspension was then prepared by diluting 0.5 µL of beads in 360 µL of water, added directly to the MgCl₂ solution present in the chamber, and incubated for a further 3 min. The liquid was then removed and replaced with water to wash away non-adherent beads. Regions containing well-isolated, properly immobilized beads were then selected under the microscope. For each field of view, a z-stack was acquired from –1 µm to +1 µm with a 20 nm step size, centered on the focal plane corresponding to the bead’s center and using the 640 nm laser at 1 mW and an exposure time of 100 ms. Between 7 and 10 fields of view, each containing 5 to 10 beads, were acquired per calibration.

#### Microscopy

All data were acquired on a custom microscope^25^ at room temperature. For dual-color 3D imaging in fixed cells we employed a dual-channel detection unit^25^ that splits the emission with an imaging grade long pass dichroic 665 (Chroma) and 700/100 nm and 676/37 nm bandpass filters. For biplane imaging a 50:50 non-polarizing beam splitter cube (Thorlabs) was used to split the channels, and the focusing lens in one channel was moved to create two focal planes on one camera chip. During imaging, the UV laser pulse width was automatically adjusted to keep a constant density of localizations. Exposure times were 25 ms.

Localizations were drift-corrected (**Figure 2A,B**, DECODE-Plex/GlobLoc) and optionally debiased (**Figure 2A**, DECODE-Plex) and filtered on localization precision below 30 nm.

## Data availability

The data generated and analyzed in this study are available from the Zenodo repository at https://doi.org/10.5281/zenodo.20088009. The repository includes the trained network weights, raw calibration bead images with corresponding calibrated PSF files, all datasets used for evaluation in the manuscript, and the resulting localization outputs and analysis files required to reproduce the reported results.

## Code availability

The source code for DECODE-Plex is available at GitHub (https://github.com/ries-lab/DECODE-Plex). The repository contains Jupyter notebooks that enable step-by-step reproduction of the analyses presented in this manuscript. Instructions for installation, configuration and execution are provided in the repository.

## Acknowledgements

This work was supported by the Austrian Science Fund (FWF 10.55776/ESP3754525 to Z.M.), the Bundesministerium für Bildung und Forschung (BMBF FKZ 01IS20019 ‘SiMaLeSAM’ to J.R., J.H.M. and L.R.M.), the Human Frontiers Science Program (HFSP, RGP006/2023 to J.R. and M.M.) and the European Research Council (ERC CoG724489 to J.R.). H.S. acknowledges financial support from the China Scholarship Council (CSC) for his doctoral research in Austria. We thank Yuan Jiang and Dr. Tianlun Zhang for testing this software.

## Supplementary Figures

**Supplementary Figure 1.**
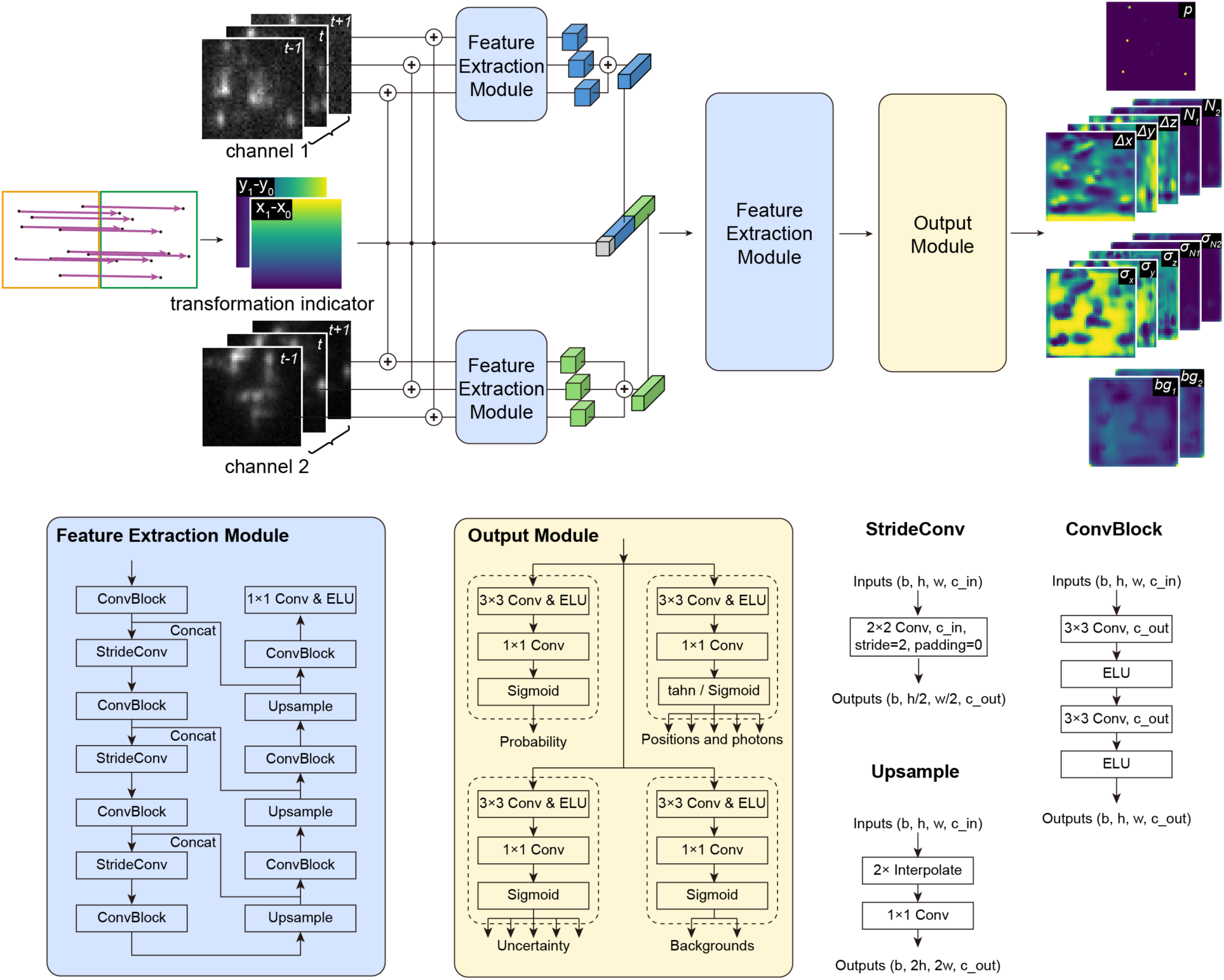
Network architecture: DECODE-Plex’s neural network backbone consists of three stages: One per-channel stage, one union stage applied on the concatenated feature maps, and one output stage producing the predictions. The per-channel stage is composed of a U-Net architecture (expanded at the bottom) and is applied to the prediction and context frames separately. Each channel has its own U-Net. In the figure, we show the case for a dual-channel scenario. The union stage is a U-Net architecture applied to the concatenation of the resulting feature maps for the prediction and context frames of all channels. To account for spatial transformations between channels, a transformation field containing the pairwise channel offsets is provided as auxiliary input. The resulting feature maps from all channels, together with the transformation field, are concatenated and processed by the joint feature extraction stage. The output stage is divided into different prediction heads, each composed of two convolutional layers. Three output heads produce probability, localization, and uncertainty estimates. An optional fourth output head can produce a background prediction. All U-Nets have three up- and down-sampling stages. Each ConvBlock consists of two 3 × 3 convolutional layers followed by ELU activations. The resolution is halved and the number of filters is doubled in each down- sampling stage, and vice versa in the up-sampling stage.

**Supplementary Figure 2.**
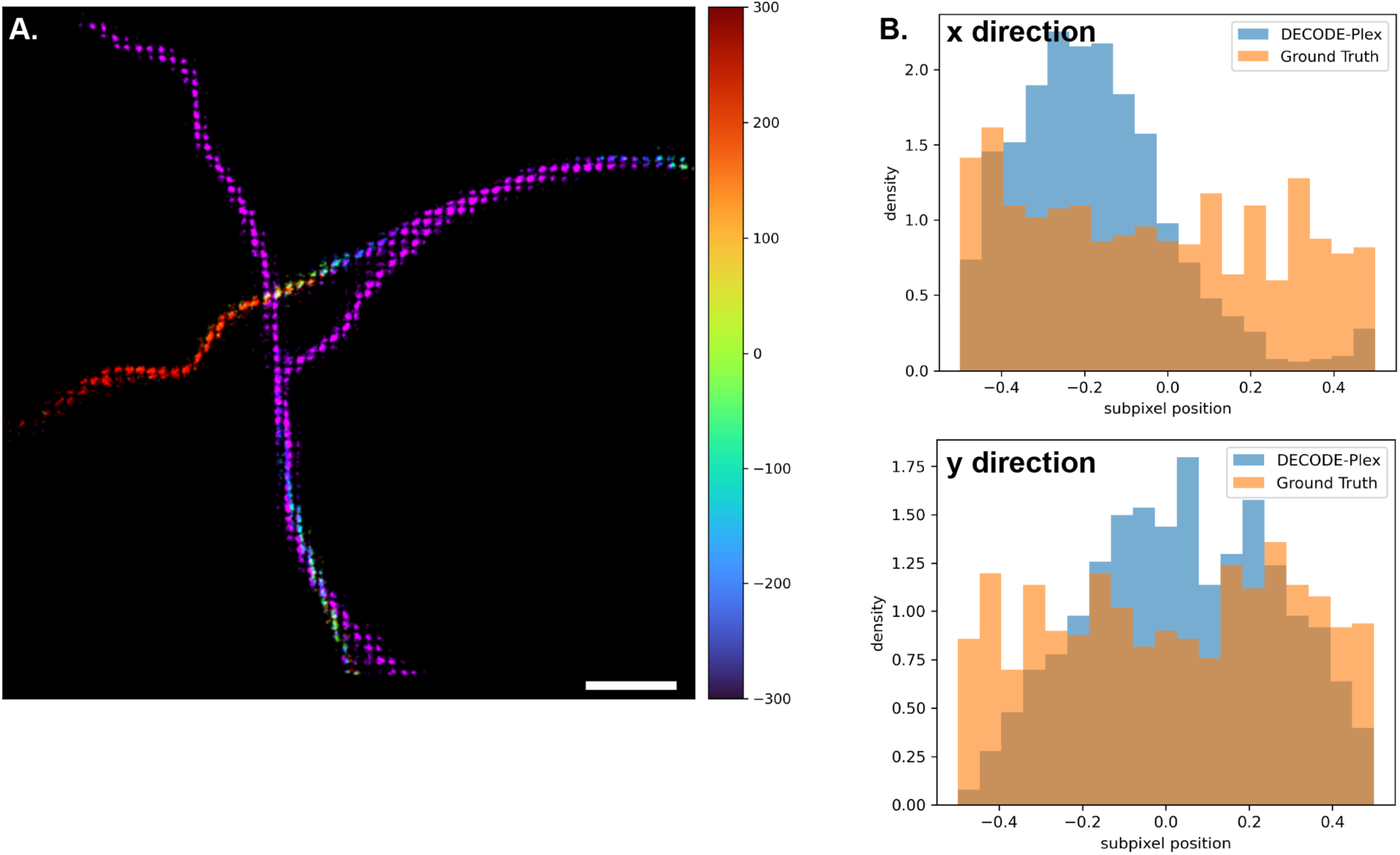
PSF model mismatch leads to pixelation. **A.** The network was trained with an incorrect PSF model and applied to the SMLM Challenge data set MT0.N2.HD-BP. DECODE-Plex then outputs gridded localizations. **B.** Subpixel positions for ground-truth emitters and matched DECODE-Plex localizations. DECODE-Plex localizations are non-uniformly distributed across the subpixel position whereas the matched ground-truth localizations are uniformly distributed. DECODE-Plex localizations are biased relative to the pixel center.

**Supplementary Figure 3.**
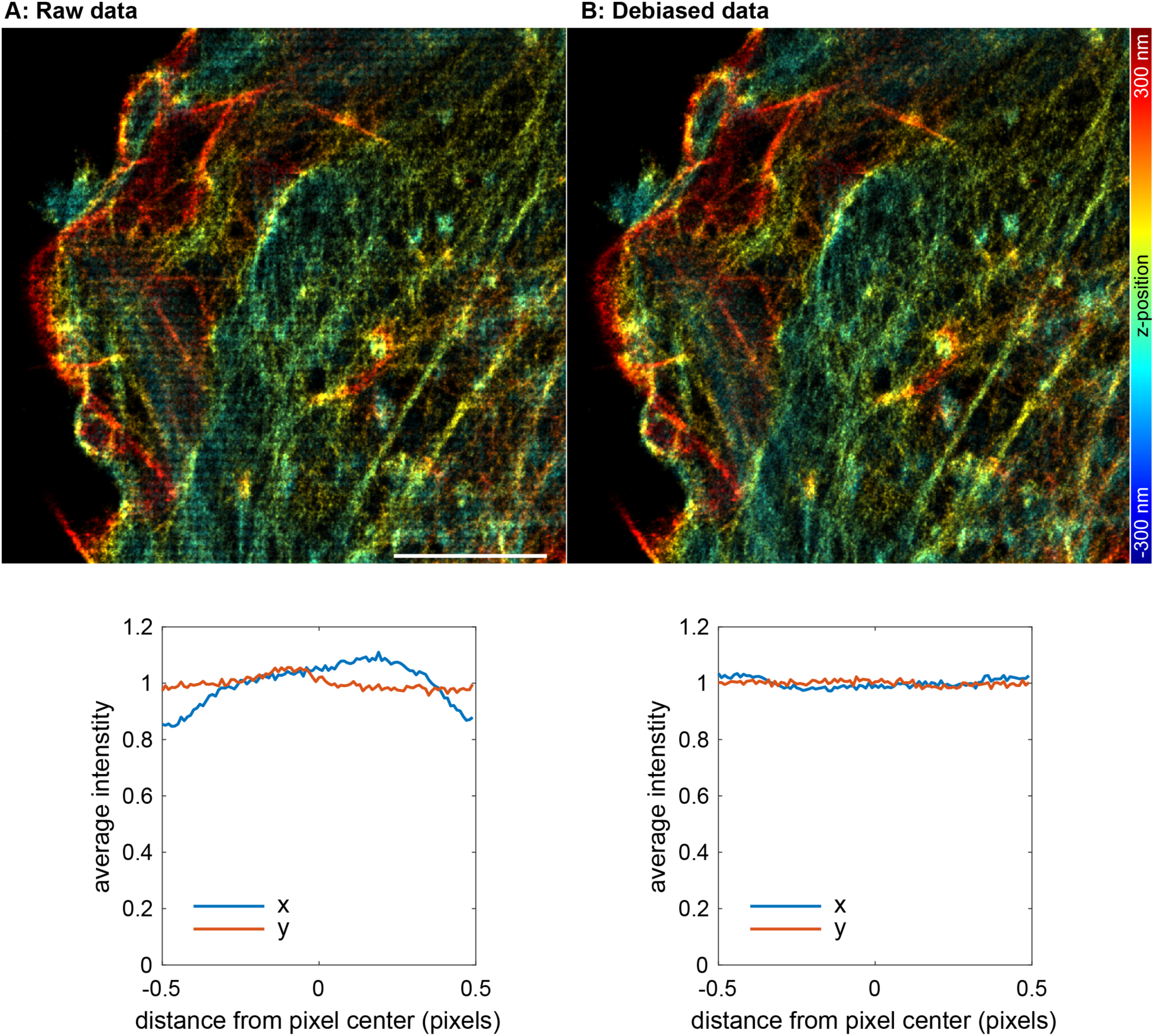
**Pixelation artifacts** on experimental data corresponding to Figure 2A (microtubules and actin). **A.** Dense, out-of-focus localizations show a gridding effect, i.e., a bias towards the pixel center. **B.** Filtered and de-biased localizations. De-biasing, which actively shifts the localizations to equalize the subpixel position histogram, reduces the pixelation artifacts.

**Supplementary Figure 4.**
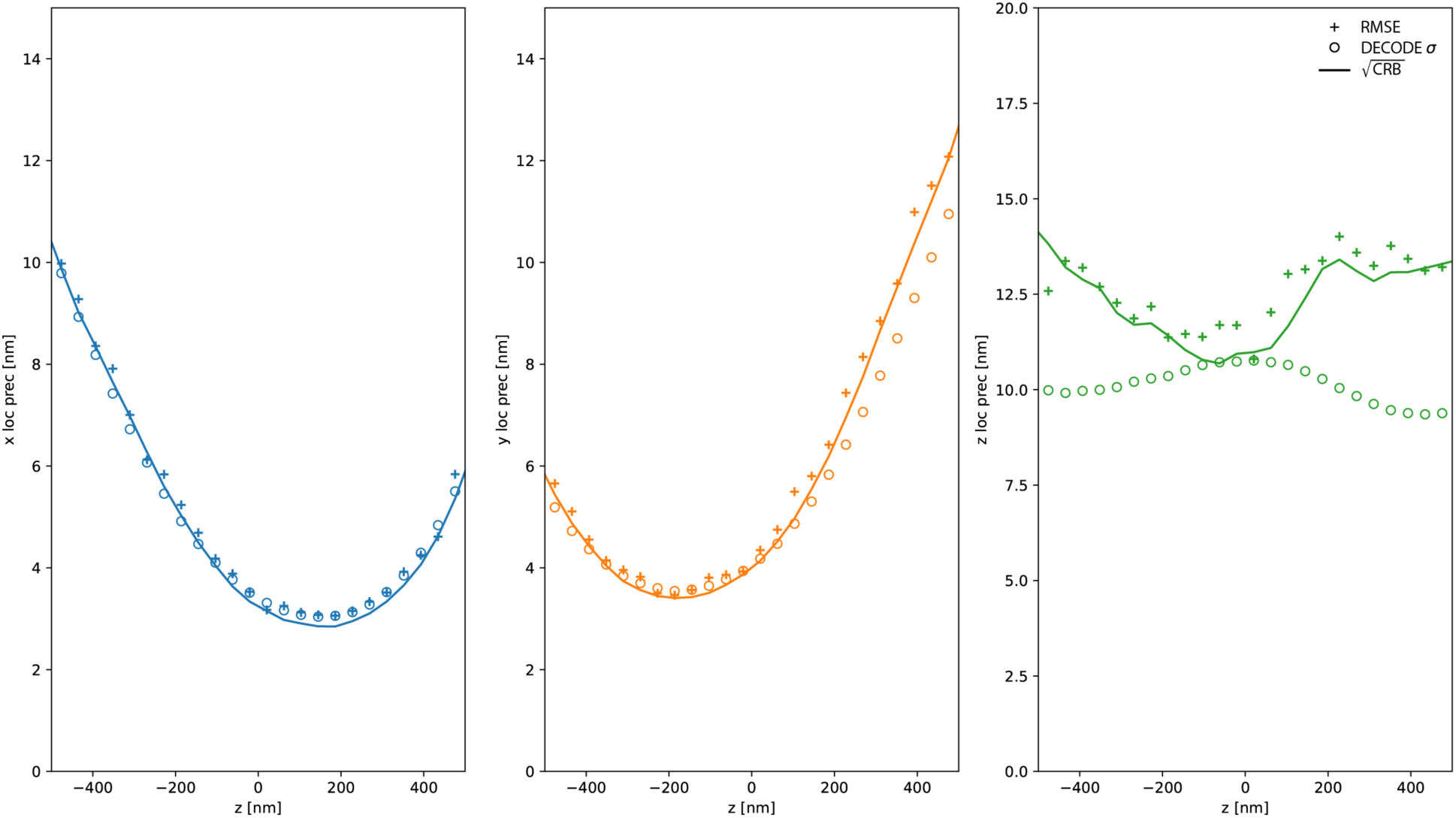
Single-emitter performance. Localization error and Cramér–Rao Bound (CRB) for dual-channel fitting. The empirical RMSE values achieved by DECODE-Plex closely match the CRB across the z-range. DECODE-Plex uncertainty estimates closely resemble the CRB in both lateral dimensions. In the axial dimension, the uncertainty values are slightly underreported. An experimental PSF and transformation were used for simulation; photons were linked, i.e., a fixed ratio of photons in the two channels was used for training.

**Supplementary Figure 5:**
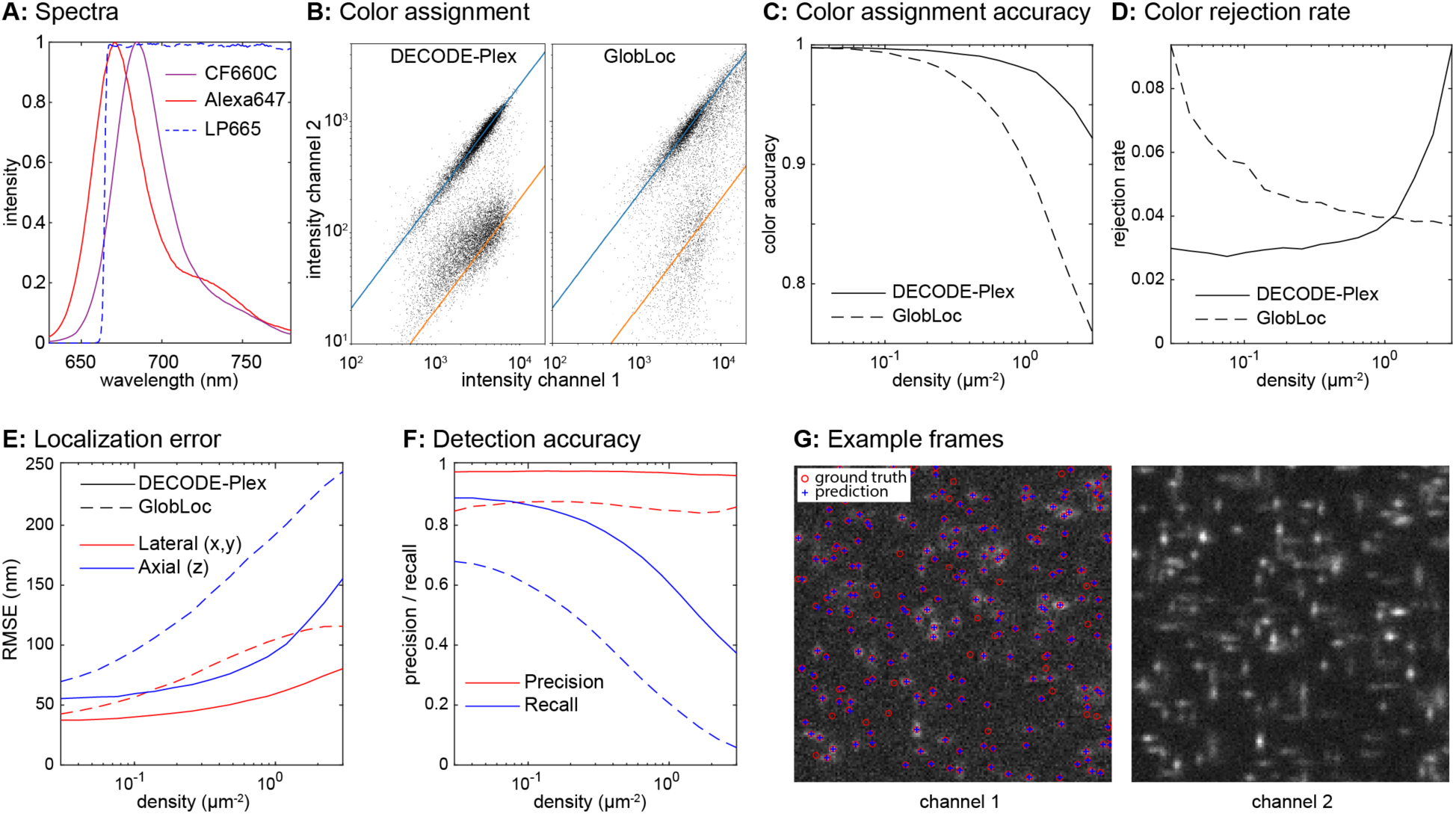
High-density dual-color fitting, performance on simulated data. **A. Spectra** of dyes and dichroic used for the simulations. **B. Color assignment** using relative intensities of both channels done with DECODE-Plex and GlobLoc. **C-F Performance metrics** (**C,** color assignment accuracy, **D,** color rejection rate, **E,** localization error, **F,** detection accuracy) comparing DECODE and GlobLoc. DECODE-Plex’s predictions were filtered for a probability threshold p < 0.5, GlobLoc’s localizations were filtered for a relative log-likehood LLrel > −1 and a photon count of phot ≥ 100. **G. Example frames.**

**Supplementary Figure 6.**
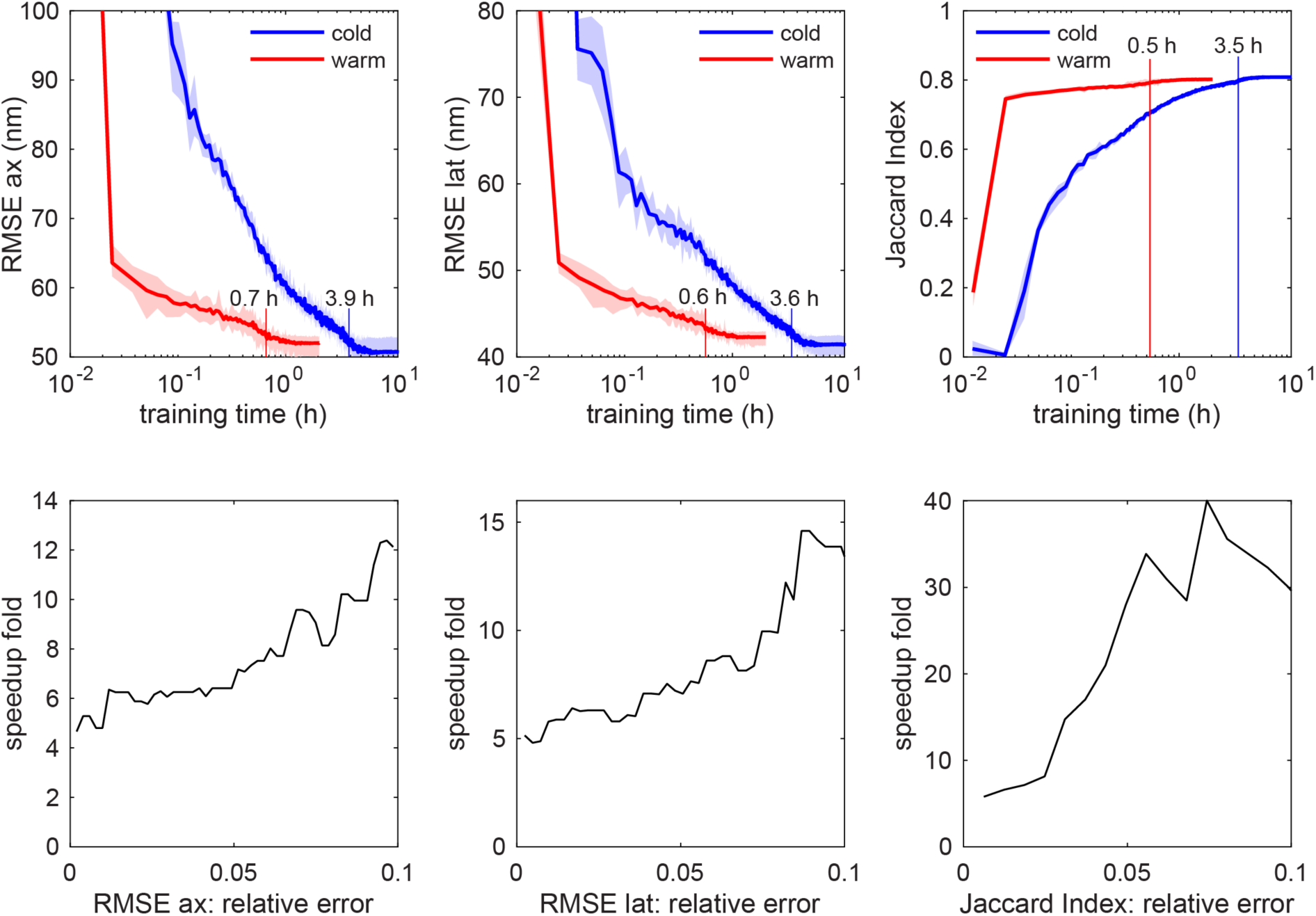
Warm start reduces training times. A base model was trained with an experimental PSF model (channel 1: 50 – 50 000 photons, 0 – 500 background photons, channel 2: 100 – 500 000 photons, 0 – 1000 background photons). A model with a different experimental PSF and different photon and background ranges (channel 1: 200 – 20 000 photons, 0 – 100 background photons, channel 2: 1000 – 100 000 photons, 0 – 500 background photons) was either trained from scratch (cold start) or initialized with a base model (warm start). Shown are axial and lateral precision (RMSE), and detection accuracy (Jaccard index) vs the training time. Shaded regions indicate minimum and maximum values from 5 (cold start) or 10 (warm start) independent trainings. The training times indicated in the graphs denote the training time after which the RMSE axial (ax) and lateral (lat) resolution was within 1 nm of the fully converged value and the Jaccard index was within 1% of the final value. The second row shows the speed up in the training for different target accuracies.

**Supplementary Figure 7.**
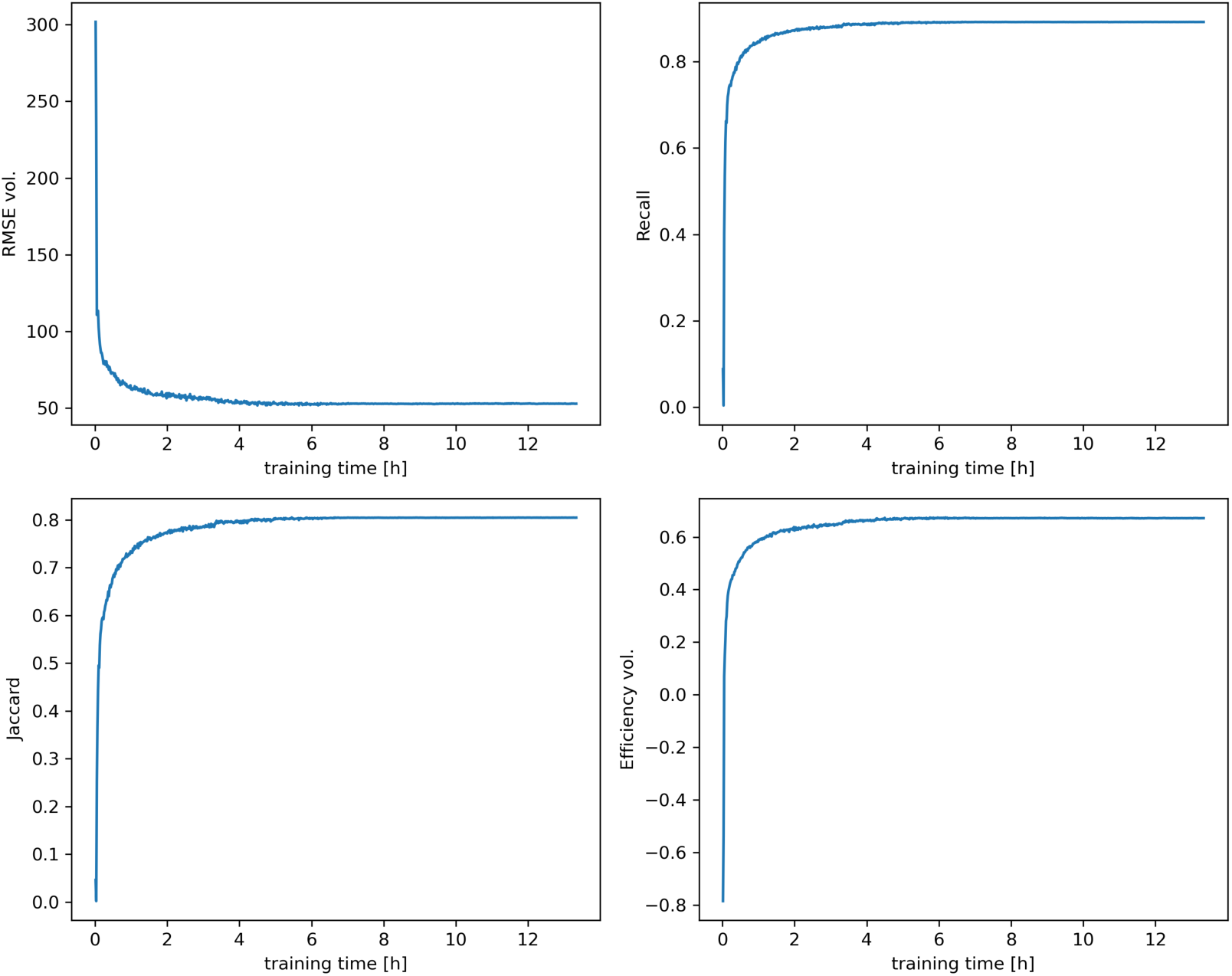
**Training convergence** as a function of the training time. Convergence of DECODE-Plex performance for several metrics. Training times are measured on a single Nvidia RTX 4090 GPU. The training data covers a common multicolor scenario at high signal-to-noise ratio.

**Supplementary Figure 8.**
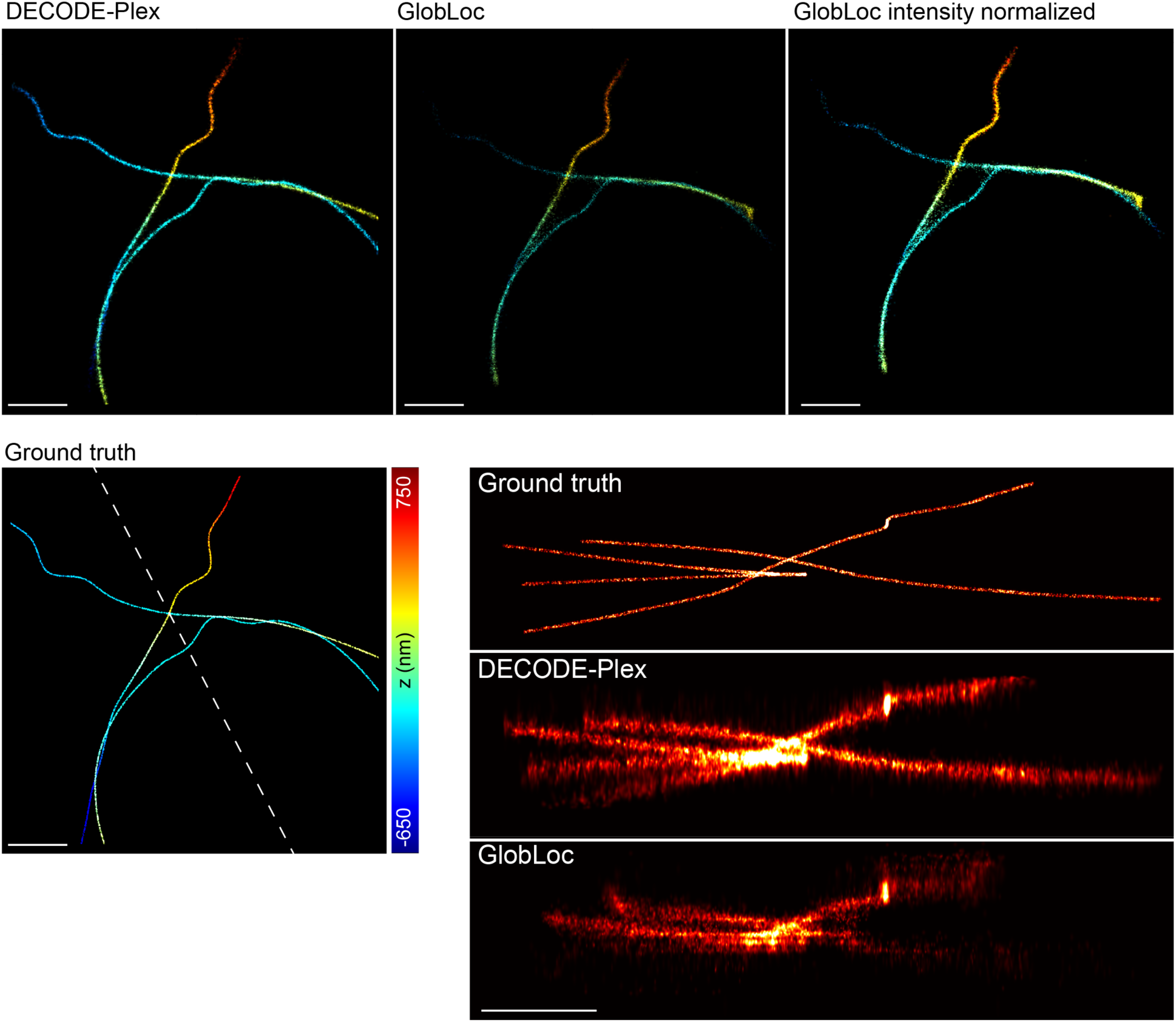
SMLM Challenge. Top-view and side-view reconstructions of the MT0.N2.HD-BP training data set. The dashed line represents the horizontal axis of the side-view reconstructions. Scale bars 1 µm.

**Supplementary Figure 9.**
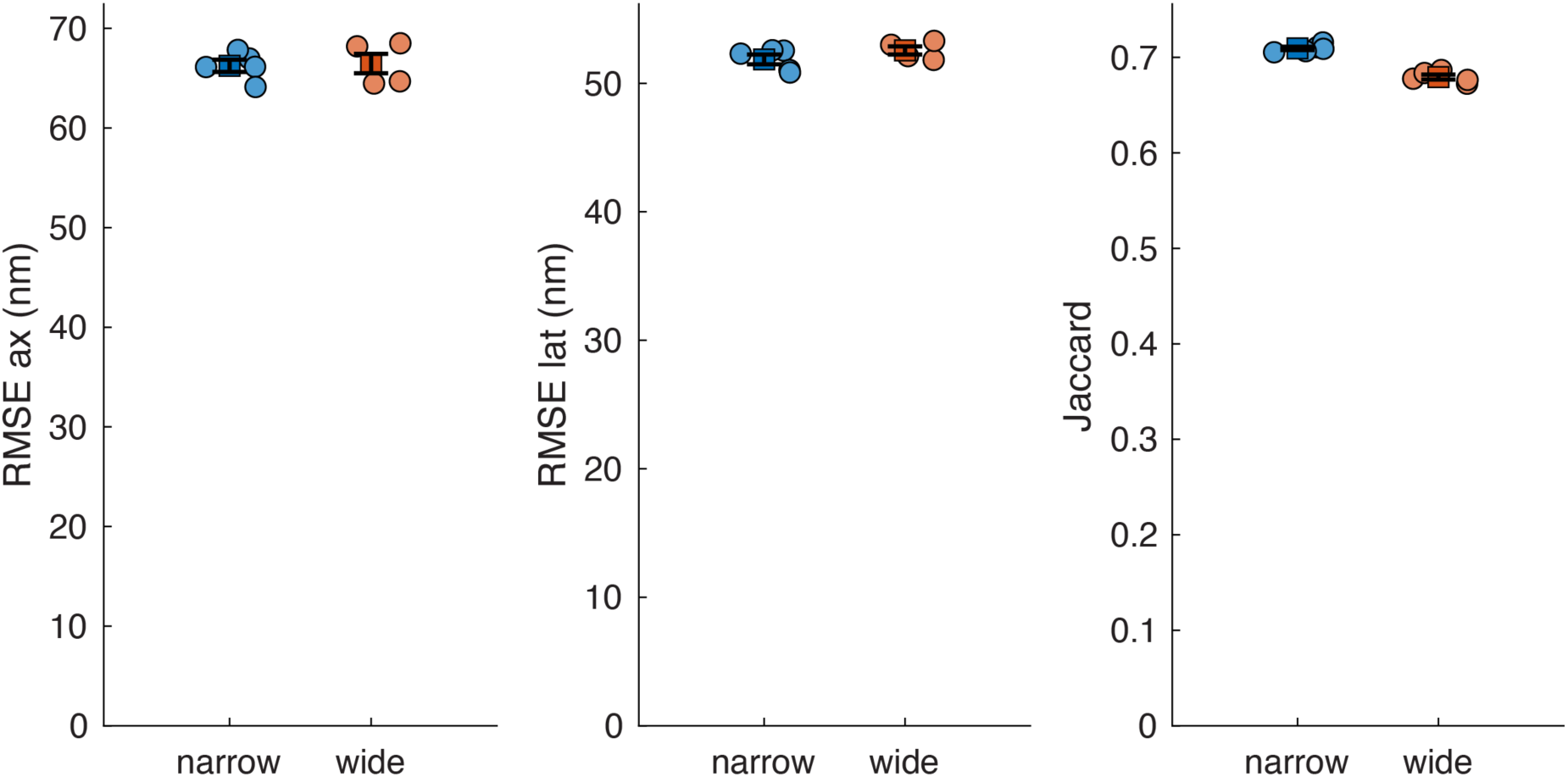
Robustness of inference with respect to training parameters. We trained network two times (narrow and wide parameter ranges) with an identical PSF but different ranges for photons and background. We used the narrow range simulations as test data for both networks. We trained the network 5 times for each condition. Here we show RMSE values in the axial and lateral direction, the Jaccard index as a measure for the detection accuracy, and the mean +/- standard deviation of the distributions. Training with a wide parameter range only occurs a small performance penalty. Wide training range: Ch1 photons: 50 – 50 000, Ch2 photons: 100 – 100 000, Ch1 background: 0 – 200, Ch2 background: 0 – 500. Narrow training range: Ch1 photons: 100 – 10 000, Ch2 photons: 200 – 20 000, Ch1 background: 50 – 100, Ch2 background: 100 – 200

**Supplementary Table 1.** Simulation and evaluation parameters for experiments. For each emitter, we draw photon counts from a uniform distribution with the range indicated in the Photon counts column. Here, “channels linked” indicates that both channels share the same total photon count, whereas “channels free” indicates that photon counts are sampled independently for each channel. For the CRLB comparisons (**Supplementary Figure 4**), we simulated 30 000 frames. For the remaining figures, we simulated frames for each combination of data density and signal-to-noise ratio until we acquired at least 100 000 emitters and 1000 frames.

| Figure | Modality | Photon counts | Background | PSF Model | Train Density [$\mu\text{m}^{-2}$] | Test Density [$\mu\text{m}^{-2}$] |
| --- | --- | --- | --- | --- | --- | --- |
| Fig 1c,d | Biplane<br>(channels linked) | Train: 1k-10k<br>Test: 5k | Train: 0-100<br>Test: 50 | SMAP | 2.5 | 0.03-3 |
| Fig 2a | Dual-color<br>(channels free) | 0.2k-20k (ch1)<br>1k-100k (ch2) | 0-100 (ch1)<br>0-500 (ch2) | SMAP | 2.5 | - |
| Fig 2b | Dual-color<br>(channels free) | 0.2k-10k (ch1)<br>0.4k-100k (ch2) | 0-30 (ch1)<br>50-200 (ch2) | uiPSF | 2.5 | - |
| Fig 2c | Biplane<br>(channels linked) | 0.5k-30k | 0-300 | SMAP | 2.5 | - |
| Fig S4 | Dual-color<br>(channels linked) | Train: 1k-4k<br>Test: 2.5k | Train: 0-40<br>Test: 20 | SMAP | 0.0469 | 1 emitter / frame |
| Fig S5 | Dual-color<br>(channels free) | 0.1k-10k (ch1)<br>0.1k-10k (ch2) | Train: 18-22 (ch1)<br>120-150 (ch2)<br>Test: 20 (ch1)<br>135 (ch2) | SMAP | 2.5 | 0.03-3 |

## Notes

### Competing Interest Statement

The authors have declared no competing interest.

